# Sparse input neural networks to differentiate 32 primary cancer types based on somatic point mutations

**DOI:** 10.1101/2020.05.13.092916

**Authors:** Nikolaos Dikaios

## Abstract

This paper aims to differentiate cancer types from primary tumour samples based on somatic point mutations (SPM). Primary cancer site identification is necessary to perform site-specific and potentially targeted treatment. Current methods like histopathology/lab-tests cannot accurately determine cancers origin, which results in empirical patient treatment and poor survival rates. The availability of large deoxyribonucleic-acid sequencing datasets has allowed scientists to examine the ability of SPM to classify primary cancer sites. These datasets are highly sparse since most genes will not be mutated, have low signal-to-noise ratio and are imbalanced since rare cancers have less samples. To overcome these limitations a sparse-input neural network (spinn) is suggested that projects the input data in a lower dimensional space, where the more informative genes are used for learning. To train and evaluate spinn, an extensive dataset was collected from the cancer genome atlas containing 7624 samples spanning 32 cancer types. Different sampling strategies were performed to balance the dataset but have not benefited the classifiers performance except for removing Tomek-links. This is probably due to high amount of class overlapping. Spinn consistently outperformed algorithms like extreme gradient-boosting, deep neural networks and support-vector-machines, achieving an accuracy up to 73% on independent testing data.

## 1 INTRODUCTION

THE main disciplines used for cancer diagnosis are imaging, histopathology, and lab tests. Imaging is commonly used as a screening tool for cancer and can guide biopsy in hard to reach organs to extract tissue samples for histopathological examination. Histopathology can identify cancer cells but cannot always determine the primary site where the tumour originated before metastasizing to different organs. Lab tests usually examine the presence of proteins and tumour markers for signs of cancer, but the results do not indicate the cancer location and are not conclusive as noncancerous conditions can cause similar results. Cancer cases of unknown primary receive empirical treatments and consequently have poorer response and survival rate [1]. Given that cancer is a genetic disease, genome analysis could lead to identification of primary cancer sites and more targeted treatments. Such analysis has recently become feasible due to the availability of large Deoxyribonucleic acid (DNA) sequencing datasets.

Cancer type identification using genome analysis involves gene expression signatures, DNA methylation and genetic aberrations. Gene expression might be the outcome of an altered or unaltered biological process or pathogenic medical condition and have been used as predictors of cancer types [2–6]. Abnormal DNA methylation profiles are present in all types of cancer and have also recently been used to identify cancer types [7,8]. This work focuses on a type of genetic aberration, namely somatic point mutations (SPM) which possess an important role in tumour creation.

Spontaneous mutations constantly take place, which accumulate in somatic cells. Most of these mutations are harmless, but others can affect cellular functions. Early mutations can lead to developmental disorders and progressive accumulation of mutations can cause cancer and aging. Somatic mutations in cancer have been studied more in depth thanks to genome sequencing, which provided an insight of mutational processes and of genes that drive cancer. Sometimes a mutation can affect a gene or a regulatory element and lead to some cells gaining preferential growth and to survival of clones of these cells. Cancer could be considered as one end-product of somatic cell evolution, which results from the clonal expansion of a single abnormal cell. Martincorena et al [9], explains how somatic mutations are connected to cancer though we don’t yet have full knowledge of how normal cells become cancer cells. Somatic point mutations have been used as classifiers of the primary cancer site [10–14]. The performance however of traditional classification algorithms is hindered by imbalances arising from rare cancer types, small sample size, noise and high data sparsity. Support vector machines (svm), classification trees, k-nearest neighbours perform well for data with complex relations, specifically for low and moderate dimensions, but are not suitable for high-dimensional problems. Neural networks with many layers (deep) according to the circuit complexity theory can efficiently fit complex multivariate functions and perform well on highly dimensional data. Shallower neural networks could in theory perform equally good but would require many hidden units [15]. Deep neural networks require large training datasets and sophisticated stochastic gradient descent algorithms to alleviate the vanishing gradient problem. Most genes however do not contain any mutation, which would affect the learning ability of neural networks. Machine learning algorithms such as k-means clustering [14], inter-class variations [16] have been used to find the discriminatory subset of genes to decrease the complexity of the problem. Identifying a discriminatory subset of genes will not necessarily resolve the problem of sparsity as most of the genes will still not contain a mutation.

This works proposes a sparse input neural network which overcomes this limitation using a sparse group lasso regularization. Its performance is validated against commonly used classifiers and extreme gradient boosted trees (XGBoost) [17]. XGBoost is based on gradient boosting machine and can represent complex data with correlated features (genes), is robust to noise, and can manage data imbalance to some extent. Different balancing strategies were applied as a pre-processing step to examine if their application would benefit the classification accuracy. To evaluate the proposed methodologies an extensive DNA sequencing database was collected from the cancer genome atlas [18]. The database consisted of 7624 samples with 22834 genes each, spanning 32 different cancer types.

## 2 THEORY

Neural networks are not well suited for high dimensional problems were the number of features *p* (e.g. *p*=22834) is high compared to the number of samples (e.g. n=7624). The dataset formulated in this work (described in the methods section) is a set of binary features categorized in 32 cancer types. The formulated database is a case of multi-class high dimensional data problem as the number of features *p* is high compared to the number of samples. Only 1974759 features (genes) from the whole dataset show sign of mutation, which means around 99% of the data is zero. Highly sparse datasets that contain many zeros (or contain incomplete data with many missing values) pose an additional problem as the learning power decreases due to lack of informative features. To predict the response of such a complex problem lasso (least absolute shrinkage and selection operator [19]) terms could be used in the objective function of the neural network to ensure sparsity within each group (cancer type) [20]. The l1 regularization of the neural network first layer weights θ, |*θ*|_1_can result in sparse models with few weights. Consequently, when p>n it is possible lasso will tend to choose only one feature out of any cluster of highly correlated feature [21]. More than one genes are commonly encoding a cancer type hence they should all be included and excluded together. This can be ensured by group lasso [22], which can result in a sparse set of groups but all the features in the group will be non-zero. A sparse group lasso penalty suggested by Simon et al (2013) [23] mixes lasso and group lasso to achieve sparsity of groups and of the features within each group, which better suits the problem at hand. An extension of the sparse group lasso [24] that groups the weights of the first layer to the same input to select a subset of features and uses an additional ridge penalty to the weights of all layers other than the first to control their magnitude was used in this work.

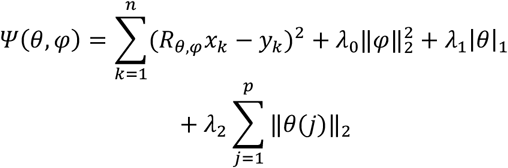

R_θ,φ_ is the network structure with θ the weights of the first input layer and φ the weights of all layers other than the first, x is the p dimensional feature (input) vector, y is the response variable and λ are the regularization parameters. *x* is a binary vector of length p = 22834, where the *i^th^* component is 0 if the *i^th^* gene is not mutated and 1 if the *i^th^* gene is mutated.

## 3 METHODS

### 3.1 Final Stage

The dataset described in the results section (22834 genes from 7624 different samples spanning 32 cancer types) was split into two sets of samples ensuring the same proportions of class labels as the input dataset: one with 90% training and 10% testing data and the other with 80% training and 20% testing data. Samples were shuffled before splitting and split in a stratified way to ensure the same proportions of class labels between the training and testing dataset. The splitting of the training and testing datasets was repeated 10 times to avoid a misrepresentation of the actual performance of the classifiers due to the particular features of the training and testing dataset in one split. Hyperparameters and/or model parameters were optimized using a grid search approach for each classifier as part of the training. The optimal were selected based on the best mean cross validation accuracy. Ten-fold cross validation was performed for all algorithms. Machine learning algorithms were developed in Python written in Keras [25] with a Tensorflow backend [26]. The developed algorithms were decision tree, k-nearest neighbors, support vector machines, artificial deep neural network, extreme gradient boosting (xgboost) and sparse input neural nets (spinn).

k-nearest neighbors run with k=5. Decision trees run with maximum depth of the tree equal to 50 and minimum number of samples required to split an internal node equal to 20. Support vector machines run with regularization parameter C=0.1 and kernel coefficient gamma=100.

Deep neural network run with 4 hidden layers with 8000 neurons each, total training epoch 70; learning rate [0.001, 0.01, 0.1, 0.2]; weight decay 0.0005; and the training batch 256. The Relu activation function was used, and the softmax at the final layer. This a multi-class classification problem where the labels (cancer types) are represented as integers hence a sparse categorical cross entropy objective function was used instead of a categorical cross entropy. Given the relatively large number of classes (i.e. 32) softmax function would be quite slow to calculate among all of them in case a categorical cross-entropy was used. Sparse categorical cross entropy only uses a portion of all classes significantly decreasing the computational time. Training was performed by minimizing the sparse categorical cross-entropy, using adaptive learning rate optimization (ADAM [27]).

XGBoost is a fast implementation of a gradient boosted decision tree. Decision trees are regression models in the form of a tree structure, however they are prone to bias and overfitting. Boosting is a method of sequentially training weak classifiers (decision trees) to produce a strong classifier where each classifier tries to correct its predecessor to prevent bias-related errors. Gradient boosting tries to fit the errors made in the initial fit and correct the corresponding errors in further training. XGBoost run with maximum tree depth 12, boosting learning rate 0.1 number of boosted trees to fit 1000, subsampling parameter in 0.9 and sampling level of columns by tree 0.8. A softmax objective function was used, and multiclass log-loss as an evaluation metric.

Spinn (described in the theory section) run with 3 hidden layers (with 2000, 1000, 500 neurons respectively), maximum number of iterations 1000, λ0=0.0003, λ1=0.001 and λ2=0.1. Training was performed by minimizing the objective function at equation 1 using ADAM [27].

### 3.2 Sampling strategies

The two main strategies to deal with imbalanced datasets is either to balance the distribution of classes at the data level or to change the classifier to adapt to imbalance data at algorithmic level. Data level balancing can be achieved by under-sampling, over-sampling or combination of both. The sampling strategies examined in this work were:

1. Random over-sampling where a subset from minority samples is randomly chosen, these selected samples are replicated and added to the original set
2. Synthetic Minority Over-sampling Technique (SMOTE) [28,29] where oversampling of minority class is done by generating synthetic samples
3. Adaptive Synthetic (ADASYN) [30] uses the distribution of the minority class to adaptively generate synthetic samples
4. Random under-sampling randomly removes data from the majority class to enforce balancing.
5. Tomek link [31] was published as a modification on CNN (Condensed Nearest Neighbour) [32] and removes samples from the boundaries of different classes to reduce misclassification
6. One-sided selection (OSS) [33] where all majority class examples that are at the boundary or is a noise were removed from the dataset
7. Edited nearest neighbour (ENN) [34] remove a sample from the class when the majority of its *k* nearest neighbours correspond to a different class
8. Combination of over and under-sampling, which was performed using smote with Tomek-links and smote with edited nearest neighbours.

## 4 RESULTS

### 4.1 Reported dataset

The dataset was collected from TCGA (The Cancer Genome Atlas) [18] with filter criteria IlluminaGA_DNASeq_Curated and was last updated on March 2019. All data can be found at http://tcga-data.nci.nih.gov. It contains information about somatic point mutations in 22834 genes from 7624 different samples with 32 cancer types. The inclusion of 32 tumour types and subtypes increases the number of associations between tumours and the number of convergent/divergent molecular subtypes.

The cancer types are abbreviated as: Adrenocortical carcinoma (ACC), Bladder Urothelial carcinoma (BLCA), Breast Invasive Carcinoma (BRCA), Cervical squamous cell carcinoma and endocervical adenomacarcinoma (CESC), Cholangiocarcinoma (CHOL), Colon adenocarcinoma (COAD), Lymphoid Neoplasm Diffuse Large B-cell Lymphoma (DLBC), Esophageal carcinoma (ESCA), Glioblastoma multiforme (GBM), Head and Neck squamous cell carcinoma (HNSC), Kidney Chromophobe (KICH), Kidney renal papillary cell carcinoma (KIRP), Acute Myeloid Leukemia (LAML), Brain lower grade glioma (LGG), Liver hepatocellular carcinoma (LIHC), Lung Adenocarcinoma (LUAD), Lung squamous cell carcinoma (LUSC), Mesothelioma (MESO), Ovarian serous cystadenocarcinoma (OV), Pancreatic adenocarcinoma (PAAD), Pheochromocytoma and Paraganglioma (PCPG), Prostate adenocarcinoma (PRAD), Rectum adenocarcinoma (READ), Sarcoma (SARC), Skin Cutaneous Melanoma (SKCM), Stomach adenocarcinoma (STAD), Testicular Germ Cell Tumors (TGCT), Thyroid carcinoma (THCA), Thymoma (THYM), Uterine Corpus Endometrial Carcinoma (UCEC), Uterine carcinosarcoma (UCS) and Uveal Melanoma (UVM). An overview of the mutations per cancer type (Ca) is shown in table 1. The number of samples varies heavily between different cancer types (e.g. BRCA has 993 samples whereas CHOL has only 36 samples) making the dataset highly unbalanced.

**TABLE 1.** SUMMARY OF SOMATIC POINT MUTATIONS PER CANCER TYPE (CA).

| Ca | Sample number | Missense Mutation | Nonsense mutation | Nonstop mutation | RNA | Silent | Slice site | Total Mutation |
| --- | --- | --- | --- | --- | --- | --- | --- | --- |
| ACC | 91 | 7258 | 524 | 15 | 1012 | 2799 | 415 | 12082 |
| BLCA | 130 | 25108 | 2215 | 47 | 0 | 10140 | 605 | 38174 |
| BRCA | 993 | 55063 | 4841 | 133 | 4474 | 17901 | 1561 | 83973 |
| CESC | 194 | 26606 | 2716 | 84 | 6191 | 9765 | 574 | 45936 |
| CHOL | 36 | 3307 | 316 | 0 | 343 | 1166 | 90 | 5222 |
| COAD | 423 | 163454 | 12146 | 184 | 72963 | 65033 | 9128 | 324120 |
| DLBC | 48 | 9623 | 353 | 0 | 9 | 6230 | 188 | 16403 |
| ESCA | 183 | 33179 | 2829 | 0 | 3947 | 11770 | 851 | 52576 |
| GBM | 316 | 26462 | 2230 | 15 | 6981 | 10378 | 813 | 46899 |
| HNSC | 279 | 33260 | 2686 | 44 | 0 | 12922 | 864 | 49776 |
| KICH | 66 | 1498 | 103 | 0 | 4 | 682 | 54 | 2341 |
| KIRP | 171 | 9442 | 515 | 18 | 629 | 3704 | 697 | 15007 |
| LAML | 197 | 1529 | 117 | 0 | 110 | 510 | 54 | 2320 |
| LGG | 286 | 6098 | 397 | 7 | 119 | 2304 | 331 | 9256 |
| LIHC | 373 | 32553 | 1853 | 0 | 59 | 12192 | 1238 | 47895 |
| LUAD | 230 | 47700 | 3692 | 56 | 0 | 16433 | 1558 | 69546 |
| LUSC | 548 | 162388 | 12268 | 301 | 16736 | 59021 | 11708 | 263481 |
| MESO | 83 | 2780 | 177 | 8 | 1494 | 1158 | 196 | 5840 |
| OV | 142 | 7106 | 420 | 9 | 655 | 2492 | 244 | 10926 |
| PAAD | 150 | 19899 | 1307 | 6 | 125 | 7777 | 974 | 30138 |
| PCPG | 184 | 3271 | 83 | 0 | 2 | 795 | 61 | 4212 |
| PRAD | 332 | 7816 | 433 | 12 | 246 | 2779 | 496 | 11846 |
| READ | 81 | 15899 | 1724 | 0 | 15 | 5918 | 306 | 23862 |
| SARC | 259 | 15457 | 785 | 19 | 0 | 5592 | 332 | 22185 |
| SKCM | 472 | 243677 | 15231 | 111 | 16006 | 137574 | 10522 | 423963 |
| STAD | 289 | 87092 | 4423 | 93 | 95 | 36394 | 3852 | 132196 |
| TGCT | 156 | 2121 | 123 | 0 | 254 | 878 | 52 | 3428 |
| THCA | 406 | 3000 | 150 | 0 | 2 | 1260 | 77 | 4489 |
| THYM | 121 | 15938 | 1639 | 0 | 19 | 6001 | 342 | 23939 |
| UCEC | 248 | 121440 | 13472 | 158 | 2016 | 42131 | 1942 | 181159 |
| UCS | 57 | 6003 | 612 | 16 | 0 | 1780 | 275 | 8713 |
| UVM | 80 | 1665 | 73 | 0 | 138 | 944 | 36 | 2856 |
| Total | 7624 | 1197692 | 90453 | 1336 | 134644 | 496423 | 50436 | 1974759 |

The main objectives of the formulated dataset were to compare the performance of different sampling approaches and the proposed machine learning algorithms. To gain a better insight of the formulated dataset intra- and between-class tests were performed on the original dataset before any sampling or splitting was performed. Intra class correlations were estimated (Table 2) to examine how strong samples in the same cancer class resemble each other. Aside from MESO and LAML the samples on the other cancer types were moderate, good or excellent. Correspondence analysis was performed to determine the variables response of the genes×samples data in a low dimensional space. Correspondence analysis can reveal the total picture of the relationship among genes-samples pairs that cannot be performed by pairwise analysis and was preferred over other dimension reduction methods because our data consist of categorical variables. Cumulative inertia was calculated (Figure 1) and it was estimated that 1033 dimensions retained >70% of the total inertia, which implies overlapping information between different samples.

**Fig. 1.**
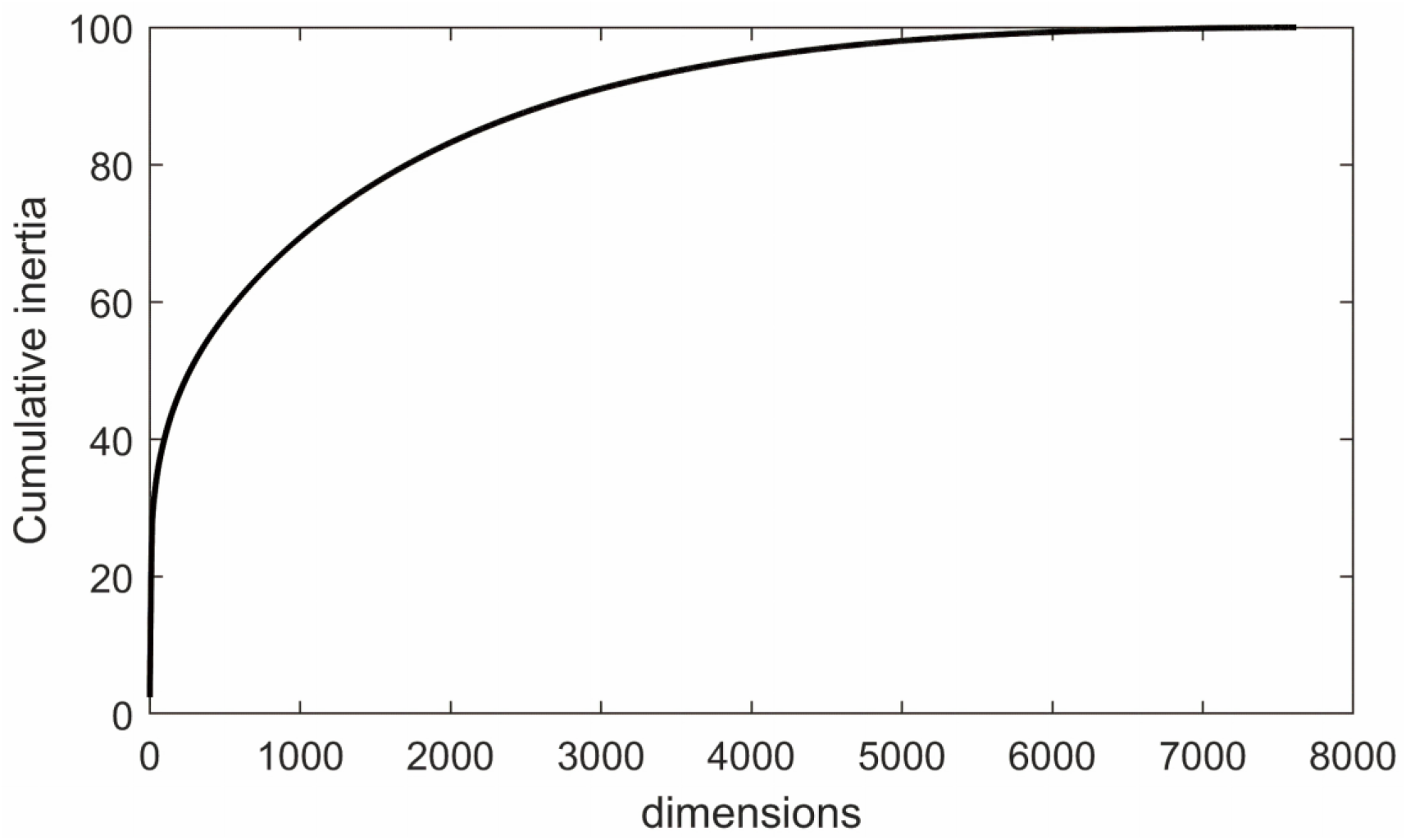
Plot of the cumulative inertia following correspondence analysis.

**TABLE 2.** INTRA CLASS CORRELATIONS FOR EACH CANCER CLASS

| ACC | BLCA | BRCA | CESC | CHOL | COAD | DLBC |
| --- | --- | --- | --- | --- | --- | --- |
| 0.526 | 0.733 | 0.888 | 0.672 | 0.854 | 0.599 | 0.914 |
| GBM | HNSC | KICH | KIRP | LAML | LGG | LIHC |
| 0.761 | 0.578 | 0.899 | 0.516 | 0.328 | 0.938 | 0.513 |
| LUSC | MESO | OV | PAAD | PCPG | PRAD | READ |
| 0.869 | 0.321 | 0.836 | 0.816 | 0.958 | 0.526 | 0.68 |
| SKCM | STAD | TGCT | THCA | THYM | UCEC | UCS |
| 0.689 | 0.648 | 0.968 | 0.548 | 0.928 | 0.623 | 0.87 |

### 4.2 Overall performance of the classifiers on the original dataset

Sparse input neural networks outperformed the other classifiers both on the 10% and 20% testing datasets (table 3). The evaluation was performed using 4 different metrics, namely accuracy, precision, recall and F-score. Accuracy (Acc) is the most commonly used metric measuring the ratio of correctly classified samples over the total number of samples, however it provides no insights on the ratio of true positives over true negatives. Precision relates to the true positive rate and equal to the ratio of true positives over the sum of true and false positives. Recall also referred as sensitivity relates to the ratio of correctly classified samples over all samples that have this cancer type and is equal to the ratio of true positives over the sum of true positives and false negatives. F-score is more complex to understand but is more reliable than accuracy in our case because the dataset is imbalanced, and the numbers of true positives and true negatives are uneven.

**TABLE 3.**
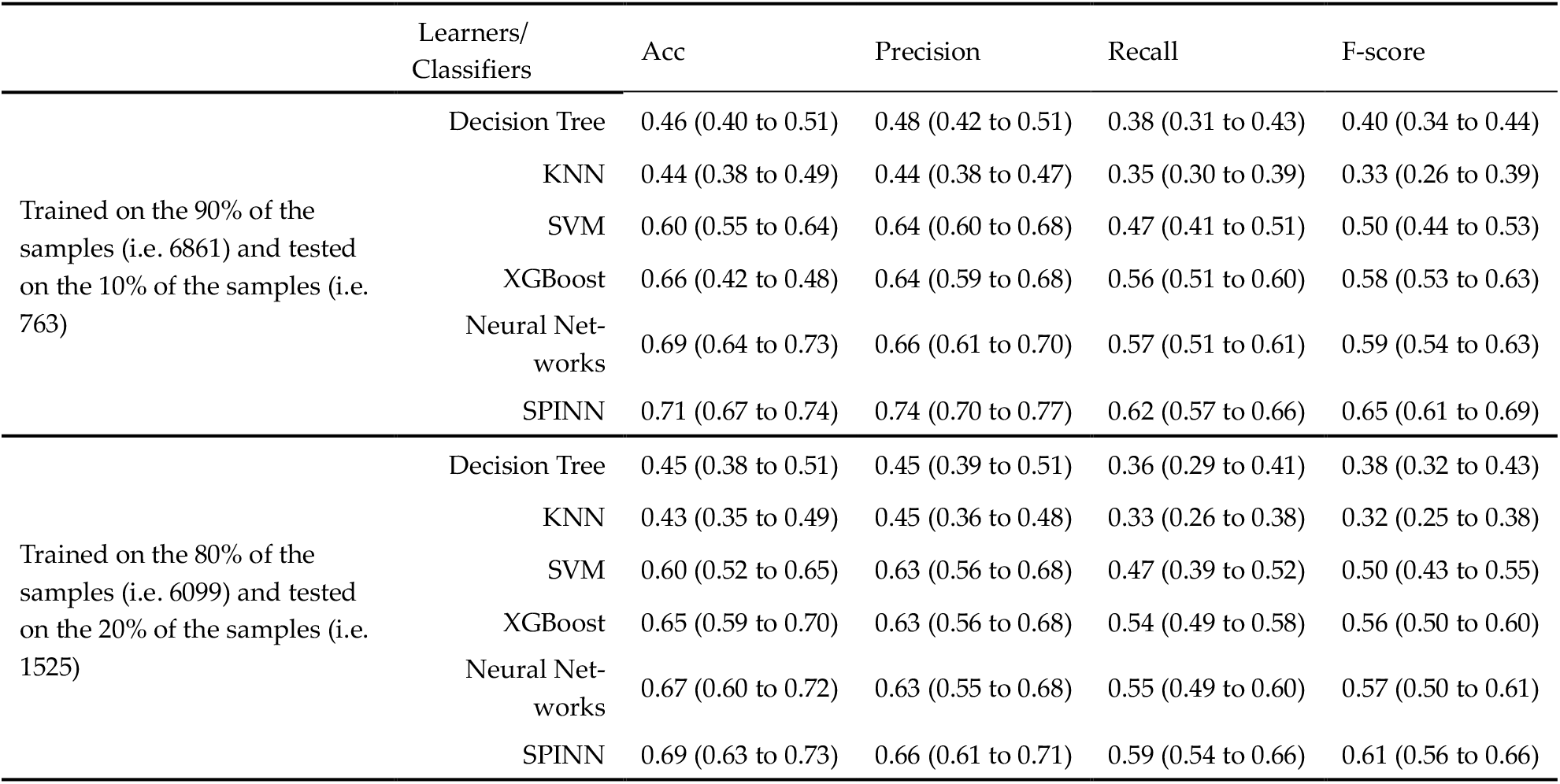
EVALUATION OF THE DIFFERENT CLASSIFIERS ON THE ORIGINAL TEST-ING DATASET. THE MEDIAN VALUES (25% TO 75% IN-TERQUARTILE RANGE) OF THE METRICS ARE REPORTED OVER THE 10 DIFFERENT SPLITS OF THE TRAINING AND TESTING DATASETS.

### 4.3 Sampling

Initially the different over/under sampling strategies were applied to the training datasets. For datasets generated after applying oversampling techniques (i.e. SMOTE and ADASYN) the performance of the classifiers remained comparable to the tests done on original datasets. SMOTE gave better results than ADASYN probably due to samples generated on the outside of the borderline of the minority class. Under-sampling methods were also applied to remove many samples from the data. In the case of ENN and CNN, the created dataset contained only 1264 and 772 samples respectively from the original data. Based on this finding one could conclude that most of the classes are overlapping and having multiple covariates, which was also implied by the correspondence analysis (Figure 1). This class overlapping can be considered as the main factor in classifiers poor performance besides class imbalance.

Due to the reduced number of samples, all classifiers performed poorly on the undersampled data. The only technique that marginally benefited classification (Table 4) was the removal of Tomek-links. This approach removes samples from the boundaries of different classes to reduce misclassification.

**TABLE 4.** EVALUATION OF THE DIFFERENT CLASSIFIERS ON THE TESTING DATASET (AFTER TOMEK-LINKS WERE REMOVED FROM THE ORIGINAL DATASET AND REDUCED TOTAL NUMBER OF SAMPLES TO 6859 FROM 7624). THE MEDIAN VALUES (25% TO 75% INTERQUARTILE RANGE) OF THE METRICS ARE REPORTED OVER THE 10 DIFFERENT SPLITS OF THE TRAINING AND TESTING DATASETS

|  | Learners/<br>Classifiers | Acc | Precision | Recall | F-score |
| --- | --- | --- | --- | --- | --- |
| Trained on the 90% of the samples<br>(i.e. 6173) and tested on the 10% of<br>the samples (i.e. 686) | Decision Tree | 0.46 (0.40 to 0.48) | 0.48 (0.42 to 0.50) | 0.38 (0.33 to 0.40) | 0.40 (0.36 to 0.42) |
|  | KNN | 0.44 (0.39 to 0.46) | 0.44 (0.40 to 0.46) | 0.35 (0.30 to 0.37) | 0.33 (0.29 to 0.36) |
|  | SVM | 0.61 (0.57 to 0.64) | 0.64 (0.60 to 0.67) | 0.47 (0.41 to 0.49) | 0.51 (0.47 to 0.53) |
|  | XGBoost | 0.68 (0.63 to 0.71) | 0.65 (0.61 to 0.67) | 0.57 (0.53 to 0.60) | 0.59 (0.56 to 0.61) |
|  | Neural Net-<br>works | 0.70 (0.65 to 0.73) | 0.65 (0.61 to 0.67) | 0.59 (0.55 to 0.63) | 0.60 (0.55 to 0.62) |
|  | SPINN | 0.73 (0.70 to 0.76) | 0.75 (0.72 to 0.78) | 0.64 (0.60 to 0.67) | 0.67 (0.64 to 0.71) |
| Trained on the 80% of the samples<br>(i.e. 5487) and tested on the 20% of<br>the samples (i.e. 1372) | Decision Tree | 0.45 (0.39 to 0.50) | 0.45 (0.40 to 0.50) | 0.36 (0.30 to 0.41) | 0.38 (0.33 to 0.42) |
|  | KNN | 0.43 (0.39 to 0.46) | 0.45 (0.40 to 0.49) | 0.33 (0.27 to 0.36) | 0.32 (0.27 to 0.35) |
|  | SVM | 0.60 (0.55 to 0.63) | 0.63 (0.59 to 0.66) | 0.47 (0.42 to 0.50) | 0.50 (0.45 to 0.53) |
|  | XGBoost | 0.66 (0.62 to 0.69) | 0.64 (0.60 to 0.67) | 0.55 (0.50 to 0.59) | 0.57 (0.51 to 0.60) |
|  | Neural Net-<br>works | 0.68 (0.63 to 0.72) | 0.66 (0.61 to 0.70) | 0.57 (0.52 to 0.61) | 0.58 (0.53 to 0.62) |
|  | SPINN | 0.71 (0.66 to 0.73) | 0.73 (0.69 to 0.76) | 0.64 (0.60 to 0.67) | 0.66 (0.61 to 0.70) |

### 4.4 Classifier performance per cancer type

Aside from the overall performance of the classifiers it is important to examine their performance per cancer type as this varies significantly. Figure 2 and 4 illustrate the performance of sparse input neural networks per cancer type (F-score) on the 10% and 20% testing dataset respectively. Figures 3 and 5 show confusion matrices on the 10% and 20% testing dataset respectively, to better understand the performance of the sparse input neural network.

**Fig. 2.**
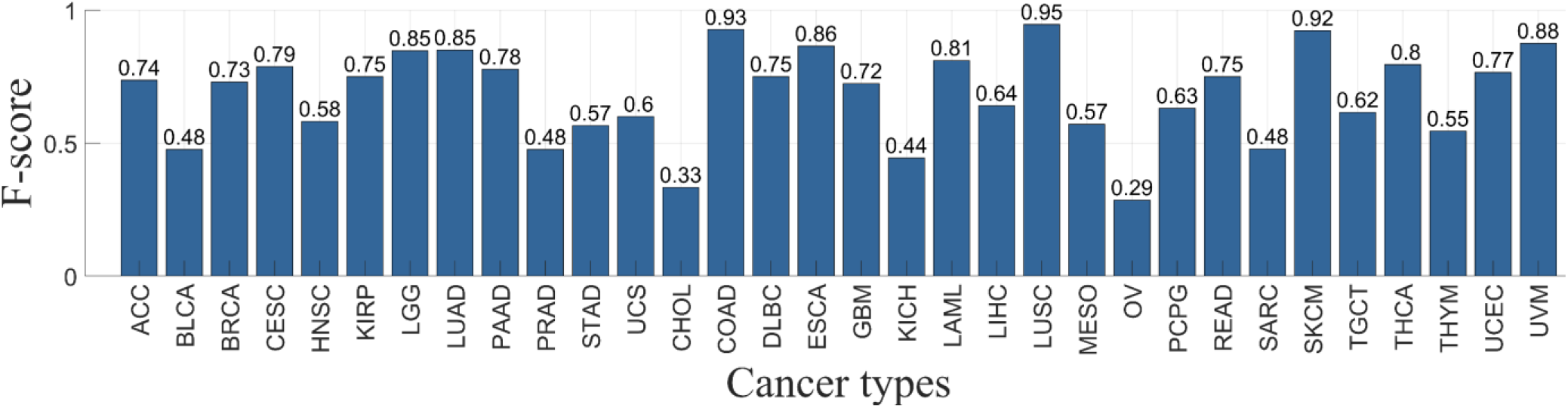
F-score (median value over 10 different splits of training and testing datasets) per cancer type for the sparse input neural network on the 10% testing dataset.

**Fig. 3.**
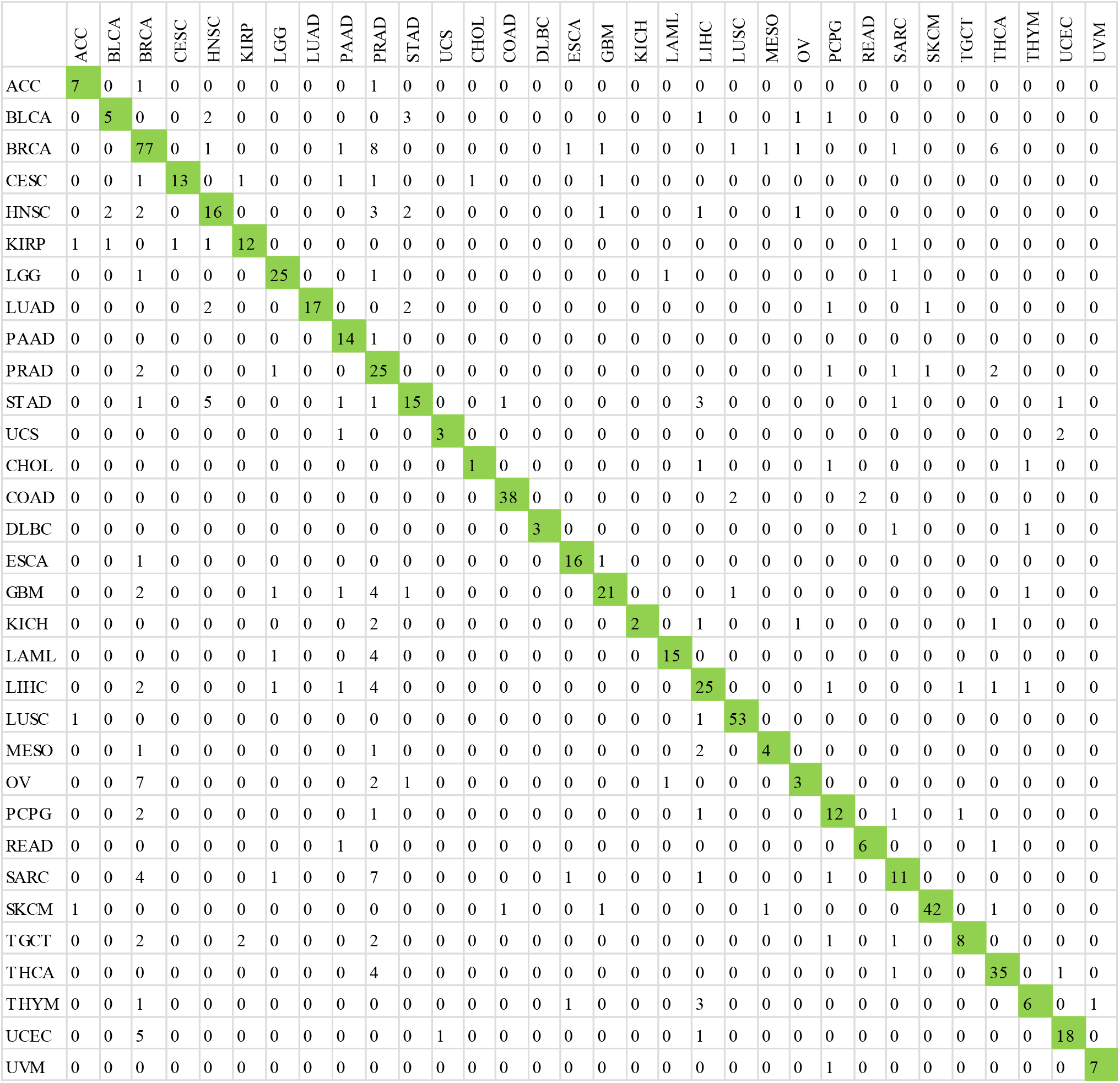
Confusion matrix of multi-class classification (columns: predicted, row: true) for the sparse input neural network on the 10% testing dataset.

**Fig. 4.**
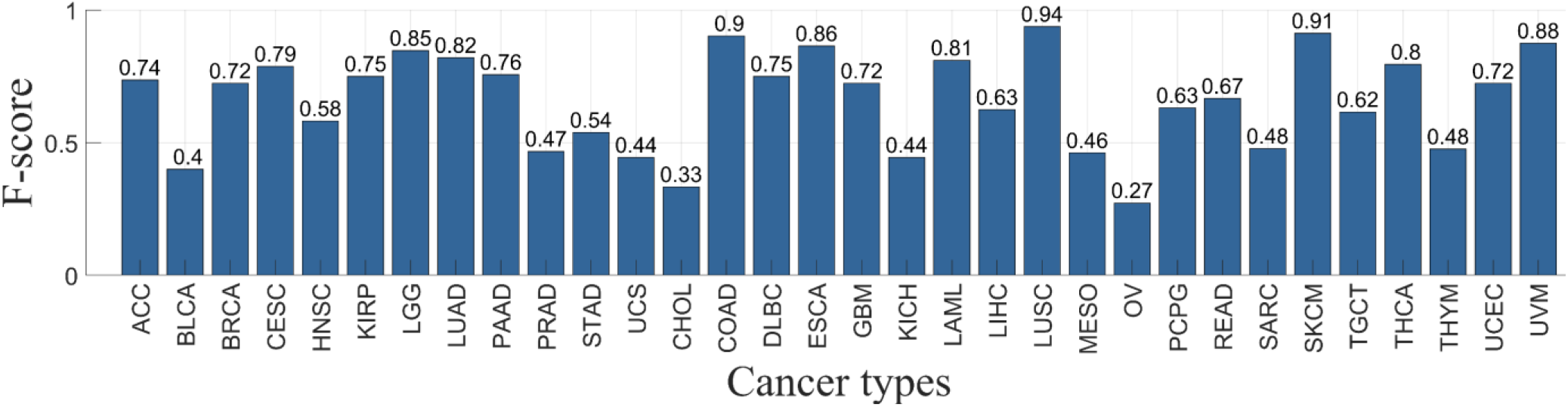
F-score (median value over 10 different splits of training and testing datasets) per cancer type for the sparse input neural network on the 20% testing dataset.

**Fig. 5.**
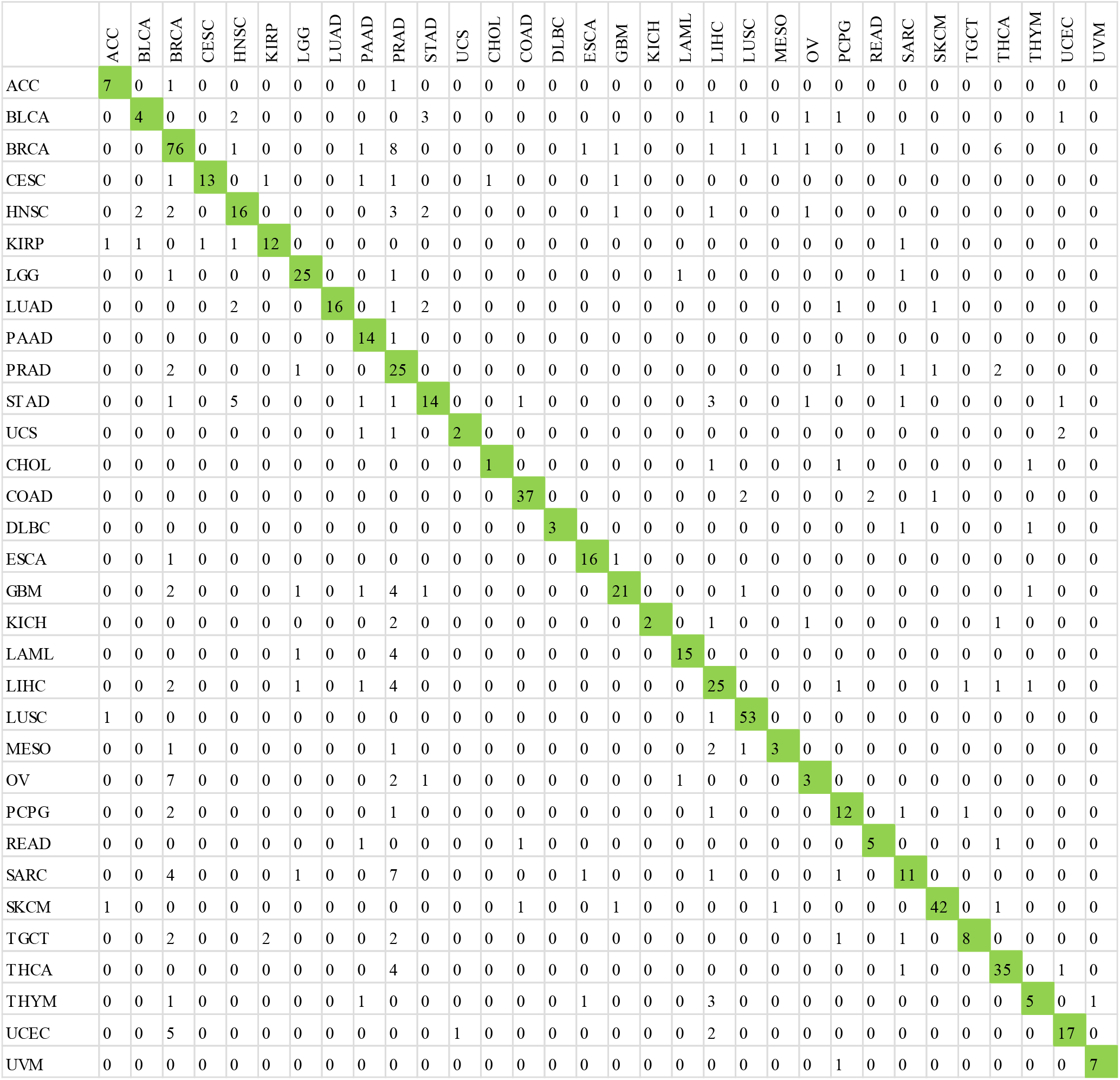
Confusion matrix of multi-class classification (columns: predicted, row: true) for the sparse input neural network on the 20% testing dataset.

As expected, the performance of classifier varies per cancer type (e.g. 0.24 for OV and 0.94 for LUSC) but this variance should not necessarily be attributed to the sample size. The Spearman's rank correlation coefficient was used to decide whether the sample number and the F-score per cancer type are correlated without assuming them to follow the normal distribution. There was no rank correlation between sample size and F-score (r=0.02 for the 10% testing dataset and r=0.04 for the 20% testing dataset).

## 5 DISCUSSION

The Cancers of unknown primary site are cancers where the site tumour originated represent ~5% of all cancer cases. Most of these cancers receive empirical chemotherapy decided by the oncologist which typically results in poor survival rates. Identification of the primary cancer site could enable a more rational cancer treatment and even targeted therapies. Given that cancer is considered a genetic disease [35], one can hypothesize that somatic point mutations could be used to locate the primary cancer type. Studies have shown promising results on identifying breast and colorectal cancer [35] but there are cancer types/subtypes where somatic point mutations are not performing well. This could be due to somatic point mutations not significantly contributing to cancer initiation but could also be a result of other limitations such as (i) high sparsity in high dimensions (ii) low signal to noise ration or (iii) a highly imbalanced dataset. As with the new cost-effective gene sequencing, we are getting a high amount of genomics data. The aim of this research is to examine the ability of somatic point mutations to classify cancer types/subtypes from primary tumour samples using state of the art machine learning algorithms.

TCGA open access data were collected as described in the methods section, which consisted of 22834 genes from 7624 different samples spanning 32 different cancer types. To the best of the authors knowledge this is the first-time such an extensive dataset with samples from 32 cancer types is reported. The resulting database is very imbalanced with common cancers sites like breast having 993 samples, while rare cancer sites having as low as 36 samples. All 22834 genes were included resulting in a highly sparse database with 99% of the genes having no mutations. Different machine learning algorithms were trained on the 90% or 80% of the original dataset and were tested on the remaining 10% or 20% respectively.

Neural networks perform well on high dimensional problems and can approximate complex multivariate functions but given that only a small subset of the genes will be informative per cancer type their performance was hindered. This work suggests a sparse input neural network (described in the theory section) which employs a combination of lasso, group lasso and ridge penalties to the loss function to project the input data in a lower dimensional space where the more informative genes are used for learning. Our results show that sparse-input neural network can achieve up to 73% accuracy on the dataset without any pre-processing of features such as gene selection. The above statement shows the learning power of neural networks with regularization. XGBoost and deep neural networks also performed well compared to traditional classifiers (decision trees, knn and svm).

All sampling strategies described in the literature are associated with the use of nearest neighbour to either oversample or undersample the dataset. In this work balancing the dataset using sampling strategies did not benefit the classifiers performance except for removing Tomek-links. This is probably due to a high amount of class overlapping. Figures 2 to 5 demonstrate that classification performance significantly varies per cancer type. In agreement with previous studies breast and colorectal cancer had a high classification accuracy (F-score up to 0.73 and 0.90 respectively). This study showcased that somatic point mutations can also accurately classify other types of cancer. There were cancer types however where classifiers performed poorly. This not necessarily related solely to having few training samples as the F-score does not seem to relate to the sample size, but for certain cancer types it could also be related to having a high amount of class over-lapping. This hypothesis was reinforced following ENN, CNN under sampling and correspondence analysis – where both suggested that only ~1000 of the samples are mutually independent.

## 6 CONCLUSIONS

To conclude this work has determined that using only somatic point mutations can yield good performance in differentiating cancer types if the sparsity of the data is considered. Results however also indicate some similarity in the information provided by somatic point mutations for different cancer types. This limitation could be managed by enriching the database especially for rare cancer types and/or introducing additional genomic information such as copy number variations, DNA methylation and gene expression signatures.

## 7 ACKNOWLEDGMENTS

This work has been supported by Royal Society Fellowship (INF\R1\191030).

**Nikolaos Dikaios** is an Assistant Professor at the Computer Vision, Speech and Signal Processig centre at the University of Surrey since 2016. He completed his DPhil (2011) in medical physics from the University of Cambridge; and worked as a research associate at University College London till 2016. His research interests are tomography, mathematical optimisation, cancer informatics and physics. Since 2019 he is a Royal Society Fellow working with Elekta on the world's first linear accelerator integrated with high field magnetic resonance imaging (MRI). He has also been awarded with the Engineering and Physical Sciences Research Council first grant to work on optimising cancer treatment with high energy proton beams. His work on machine learning for prostate cancer detection based on multi-parametric MRI has been awarded twice with the Summa Cum Laude (top 5%) and once with Magna Cum Laude (top 15%) from the flagship conference in MRI (ISMRM). He is also one of the developers of a popular open-source software for tomographic image reconstruction, STIR. STIR has been awarded the Rotblat Medal for the most cited research paper published by Physics in Medicine & Biology out of more than 3,000 articles since 2012.

